# Small Single-Stranded DNA Structure Prediction

**DOI:** 10.1101/2025.09.10.675391

**Authors:** Yutong Shi, Yu Shi, Jianing Li

**Author notes:** These authors contributed equally to this work (Yutong Shi, Yu Shi).

## Abstract

Small single-stranded DNAs (ssDNAs) shorter than 100 bases, including antisense oligonucleotides and aptamers, have been extensively studied as promising therapeutics. Their three-dimensional (3D) structures can guide development and enhance clinical potential, but obtaining these structures is often costly and time-consuming. Only a limited number of small ssDNAs have their 3D structures experimentally determined. Therefore, there is a significant need to develop accurate computational tools to predict the 3D structures of small ssDNAs. In this work, we analyze two current strategies for ssDNA structure prediction: a traditional multistep approach and the AlphaFold 3 (AF3) strategy. We compare these strategies using a dataset of 36 ssDNAs, representing 7 unique structural motifs. Our results show that AF3 can be more efficient and accurate than the traditional strategy for predicting ssDNA structures, achieving a success rate of 56% (20 out of 36 cases). However, AF3 faces some challenges when predicting long ssDNA sequences or complex motifs, such as G-quadruplexes (G4) and junctions. Interestingly, molecular dynamics (MD) simulations can significantly improve the predicted structures generated by AF3. These findings offer valuable insights into the latest ssDNA structure prediction techniques and provide a practical guide for sampling ssDNA structures with AF3 and MD.

## Introduction

Small nucleic acids have been extensively studied for their ability to modulate gene expression and target various diseases. At least 20 antisense oligonucleotides (ASOs) and aptamers have been approved as nucleic acid therapeutics for a wide range of conditions, such as amyloidosis, amyotrophic lateral sclerosis, Duchenne muscular dystrophy, etc.^1,2,3^ ASOs are synthetic DNAs or RNAs of 18 to 30 bases in length that bind to complementary mRNA molecules, to block the translation process or promote the degradation of the mRNA, thereby reducing the expression of the targeted protein.^4^ Longer than typical ASOs, aptamers are short, ssDNAs or RNAs of 20 to 100 bases, which fold into specific three-dimensional (3D) structures and function as affinity agents to interact with small molecules, proteins, or cells effectively.^5,6^ Over the years, it has been accepted that many ASOs and aptamers fold into specific, regular structures.^7^ In practice, chemical modifications are often needed to improve the druglike properties regarding stability, immunogenicity, and permeability.^8,9^ The structural knowledge of ASOs and aptamers can be useful to guide the design and selection of small nucleic acid therapeutics, as well as to impact the development of drug delivery systems.^10^ However, it can be costly and lengthy to determine the 3D structures of these nucleic acids with X-ray crystallography or nuclear magnetic resonance (NMR), while they are too small for cryogenic electron microscopy (cryo-EM).^11^ Current computational approaches can provide complementary tools to address this challenge, but they are mostly developed for RNAs, especially for larger structures.^12,13,14^ In May 2024, AF3 was released, which offers a new strategy for biomolecular structure predictions.^15^ Initial success as well as limitations have been reported in prior studies on DNA nanomotif and double-stranded DNA (dsDNA) structures,^16,17,18^ but the application to ssDNA structures has not been systematically evaluated. Therefore, we would like to fill this gap with a systematic study to develop and evaluate various approaches for accurate and effective ssDNA structure prediction, which may support the future discovery of small DNA therapeutics.

Currently, two general strategies are available for predicting small ssDNA structures **(Figure 1)**: a traditional multistep approach and a one-step direct prediction using AF3. The traditional approach starts with DNA secondary structure prediction and then constructs the 3D model, such as E2EDNA.^16^ Such an approach was built on the availability of many tools for DNA secondary structure prediction, like SeqFold^19^, SPOT-RNA^56^, UFold^57^, Mfold^58^, and MC-Fold^59^, etc. These tools implement various algorithms and energy functions, designed to predict base pairing with inputs of either DNA or RNA sequences.^19^ With the secondary structure prediction as input, tools like MacroMolecule Builder (MMB) can be used to construct the 3D structure of the target DNAs with the predicted base pairings.^20,55^ Except for a case study of a DNA-ligand complex using E2EDNA,^16^ benchmarking of the multistep traditional approach for ssDNA structure prediction has not been reported. As an alternative strategy built on the latest AI/ML advancement, AF3 can directly generate the 3D structure using the target DNA sequence as input. The training data set of AF3 includes all the data from the Protein Data Bank (PDB) available before September 2021,^15^ but dominated by B-form dsDNA entries. It remains to be assessed whether such an AI/ML approach is accurate for new ssDNA sequences or complex folding motifs. Therefore, we have conducted a systematic study to compare these two strategies and identify the need for future development of small ssDNA structure prediction. In this work, we established the traditional approach and the AF approach, followed by short MD refinement.^55^ The evaluation was conducted with 36 ssDNA structures collected from PDB, ranging from 7 to 35 bases in length, using various metrics like the *Matthews Correlation Coefficient (MCC), Global Distance Test (GDT), and Root Mean Square Deviation (RMSD)*. We have achieved good accuracy in predicting the 3D structures of short ssDNA below 14 bases in length, and MD was found to improve the prediction accuracy. However, predicting the 3D structures of long ssDNAs and complex DNA folds remains challenging, and more effective sampling and prediction approaches are needed for future applications.

**Figure 1.**
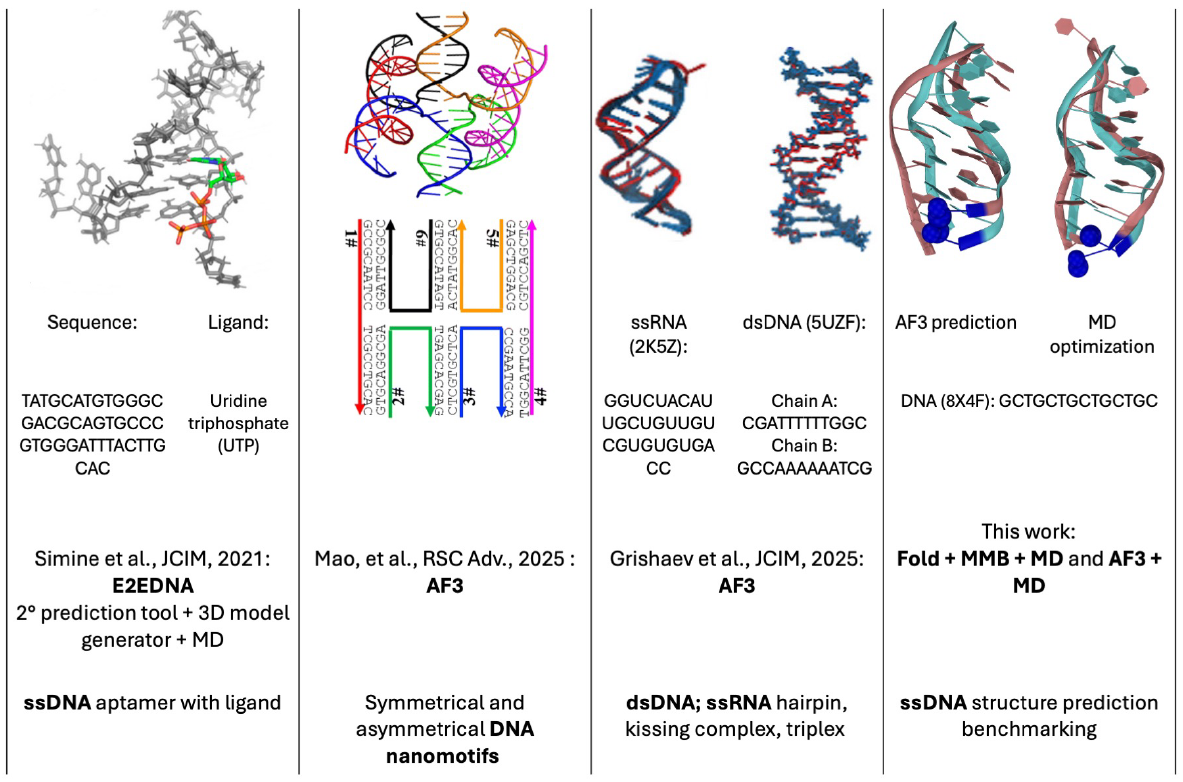
Brief overview of current DNA structure prediction studies. Previously reported approaches included E2EDNA for predicting ssDNA aptamers bound to small molecules, AF3 for DNA nanomotifs, and AF3 for dsDNA and ssRNA. This work completes the picture by focusing on ssDNA structure predictions and comparing a traditional approach (referred to as Fold/MMB) with an AI/ML approach (using AF3), along with structure refinement through all-atom MD simulations.

**Figure 2.**
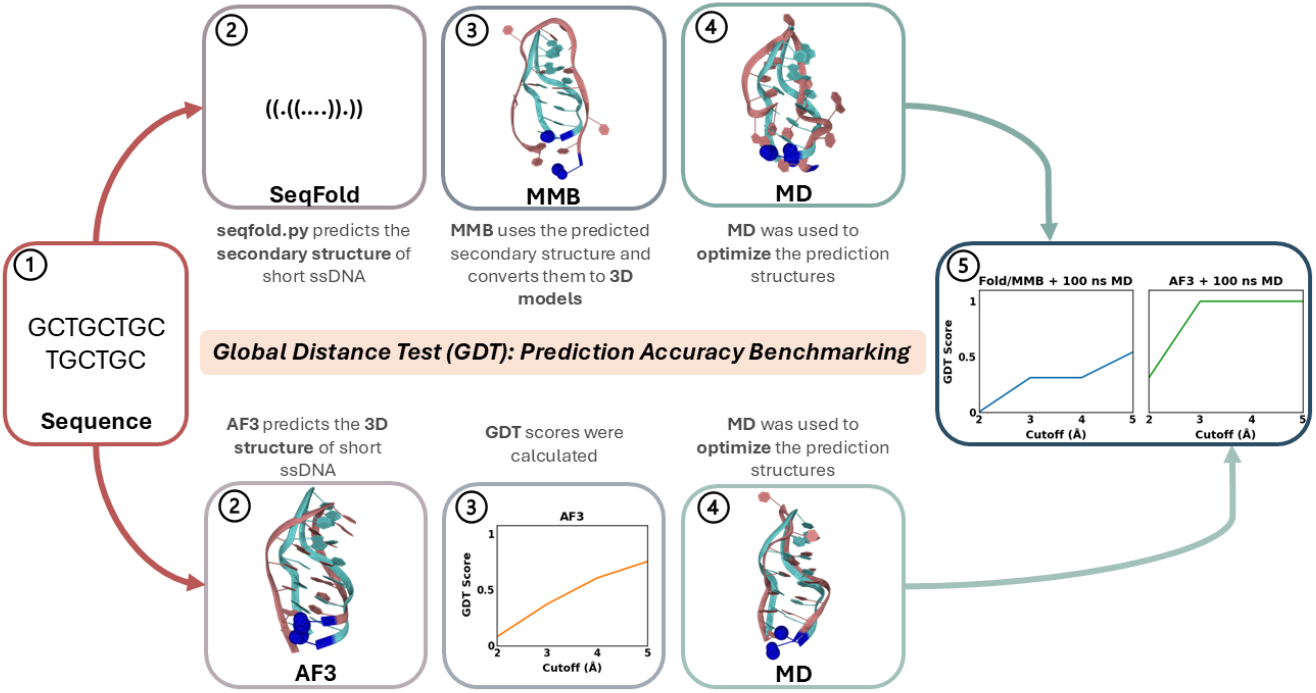
This work compares two strategies for predicting ssDNA structure. The Fold/MMB/MD pipeline (the traditional strategy) combines SeqFold, MMB, and MD, while the other pipeline (the AI/ML strategy) uses AF3 for prediction and MD for refinement.

## Models and Methods

### Data Set

We have collected 36 PDB entries of ssDNAs, which represent ssDNA folding motifs like hairpin^23,29,30,31,32,36,39,42-47,49,50,52,53,55^, short repeats^33^, minidumbbell^24,34,35,37,38^, triple helix^26^, G triplex^40,41^, G quadruplux^27,48,51^, junctions.^28,54^ Our dataset only included PDB entries without ligands, with deposit dates from 1989 to 2024. Seven representative cases, ranging from 7 to 35 bases in length, were selected for the MD optimization case study with the following criteria: for sequences shorter than 20 nucleotides, cases were chosen by increasing length in increments of 3 nucleotides. The shortest sequences within 20 – 30 and 30 – 40 bases were selected. Lastly, the longest sequence was also selected **(Figure 1)**.

### Traditional Prediction Approach

Published in 2021, E2EDNA was reported to include a two-step approach.^16^ With a target sequence, the first step was to predict the secondary structure using the SeqFold program. The secondary structure prediction was then used as the input to generate the 3D structure using the MMB program. The overall 3D structure can be optimized with MD simulations with or without ligands.^16,55^ To compare with the AI/ML approach, we adopted the traditional approach similar to E2EDNA and named it the Fold/MMB approach. We thoroughly explored the accuracy of secondary and 3D structure predictions, the impact of force fields, and the entire prediction strategy. In particular, the accuracy of secondary structure is calculated with MCC scores, and the tertiary structure accuracy is measured with GDT scores.^21,22^

### AF3-Based Approach

The AF3 webserver was used to generate the predictions. It is employed for 3D structure prediction, combined with MD refinement using two force fields, AMBER99 and OPLS5. The accuracy of the secondary structure was assessed with MCC, while the 3D structure accuracy was evaluated using GDT scores. GDT scores were calculated to reflect the improvement in prediction after MD optimization. The effects of the different force fields were also examined.

We have compared five secondary structure prediction tools SeqFold^19^, SPOT-RNA^56^, UFold^57^, Mfold^58^, and MC-Fold^59^ and chose SeqFold, as it has a DNA structure prediction program. In the Fold/MMB pipeline, SeqFold was used to predict the secondary structure of the DNA aptamers, and MMB^20^ was then used to generate the 3D model of the predicted secondary structure, in which the output from Fold/MMB only has 3D Watson-Crick base pairs but lacks additional three-dimensional folds.^16^ In the AF3 pipeline, the predictions from AF3 are compared to the native structure.^15^ In both pipelines, the predicted secondary structures were compared with the native secondary structures published on PDB, and MD was also used after 3D structure generation to optimize the predicted structures **(Figure 1). GDT** scores^21^ were recorded for structures before their MD optimization. The structure from each frame of the MD trajectory was compared to the native structure for GDT evaluations as well. The highest GDT score was selected for each MD trajectory and its corresponding DNA aptamer. AF3 structures are optimized for 10 and 100 ns.

### Definitions of GDT and MCC

**GDT**^21^ was used to measure the accuracy of the predictions. The score calculations were carried out by calculating the percentage of phosphorus atoms (P) from the predicted structure within a certain distance cutoff **(eq. 1)**. In this paper, the reported values are within 3 Å from the reference P atom, in which the reference structures are the native structures published on PDB. The alignment of the predicted structure and the native structure was performed using MDAnalysis.

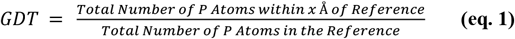

**MCC** score^22^ was used to calculate the secondary structure prediction accuracy. If the predicted pairing is in the native structure, then it is counted as a true positive (TP). If the absence of the pairing is in both predicted and native structure, it is true negative (TN). When the pairing is only shown in the predicted structure, it is false positive (FP). When the pairing is only shown in the native structure, it is a false negative (FN).^22^

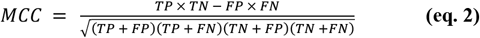

## Results

### AF3 can be efficient and accurate in predicting structures of short ssDNAs

In total, 36 ssDNA cases were predicted and analyzed using two different approaches: Fold/MMB and AF3. Of these cases, 7 AF3 predicted structures were also selected from the dataset for MD optimization, using the criteria mentioned in the method section **(Figure 3a)**. Approximately 80% of the AF3 predicted structure showed good GDT scores (0.8-1), due to the large number of short samples; shorter sequences are generally more accurate. Longer sequences showed lower GDT scores and decreased accuracy **(Figure 3)**.

**Figure 3.**
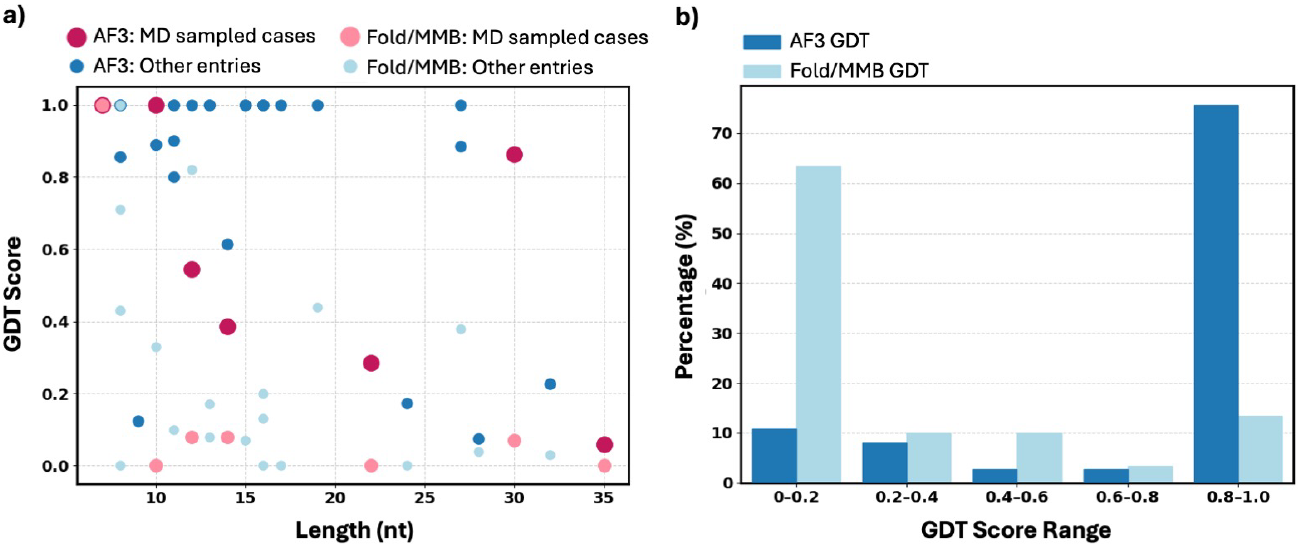
A summary of the accuracy of 36 ssDNA cases generated using Fold/MMB and AF3. a) The relationship between GDT scores and the length of the ssDNA. All other cases (dark and light blue). Selected cases for MD optimization (dark and light pink). b) Percentage of cases falls into different ranges of GDT cutoffs.

Compared to traditional secondary structure prediction tools like SeqFold, AF3 was able to generate more accurate secondary structures of the sampled aptamers. Out of the 7 cases, AF3 was able to provide 4 ssDNA – hairpin (PDBID: 1KR8), minidumbbell (PDBID: 7VM9), CTG repeats hairpin (PDBID: 8X4F), and intramolecular triplex (PDBID: 1B4Y) - with the correct secondary structures, including two cases that are not in the AF3 training regime, which consists of structures published on PDB after September 2021 – 7VM9 (released year: 2022) and 8X4F (released year: 2024). AF3 also identified 8JIC (released year: 2024) as a G quadruplex (G4) aptamer that SeqFold was unable to predict. DNAzyme (8OR8, released year: 2023) is an example of an ssDNA junction, where the SeqFold excelled with an accuracy score surpassing AF3’s accuracy, 0.70 and 0.25, respectively, and it was able to identify 8OR8 as a junction. Out of the 36 cases, SeqFold predicted 28% (10 out of 36) secondary structure correctly (MCC = 1), while 78% (28 out of 36) AF3 predicted secondary structures were correct. The secondary structures predicted by SeqFold were converted into 3D structures using MMB, and these 3D structures were compared to AF3 generated 3D structures as well. 8% (3 out of 36) cases were predicted correctly (3Å cutoff GDT = 1.00) by Fold/MMB, while 56% (20 out of 36) of the AF3 predicted structures were correct **(Table 1, S2)**.

**Table 1.**
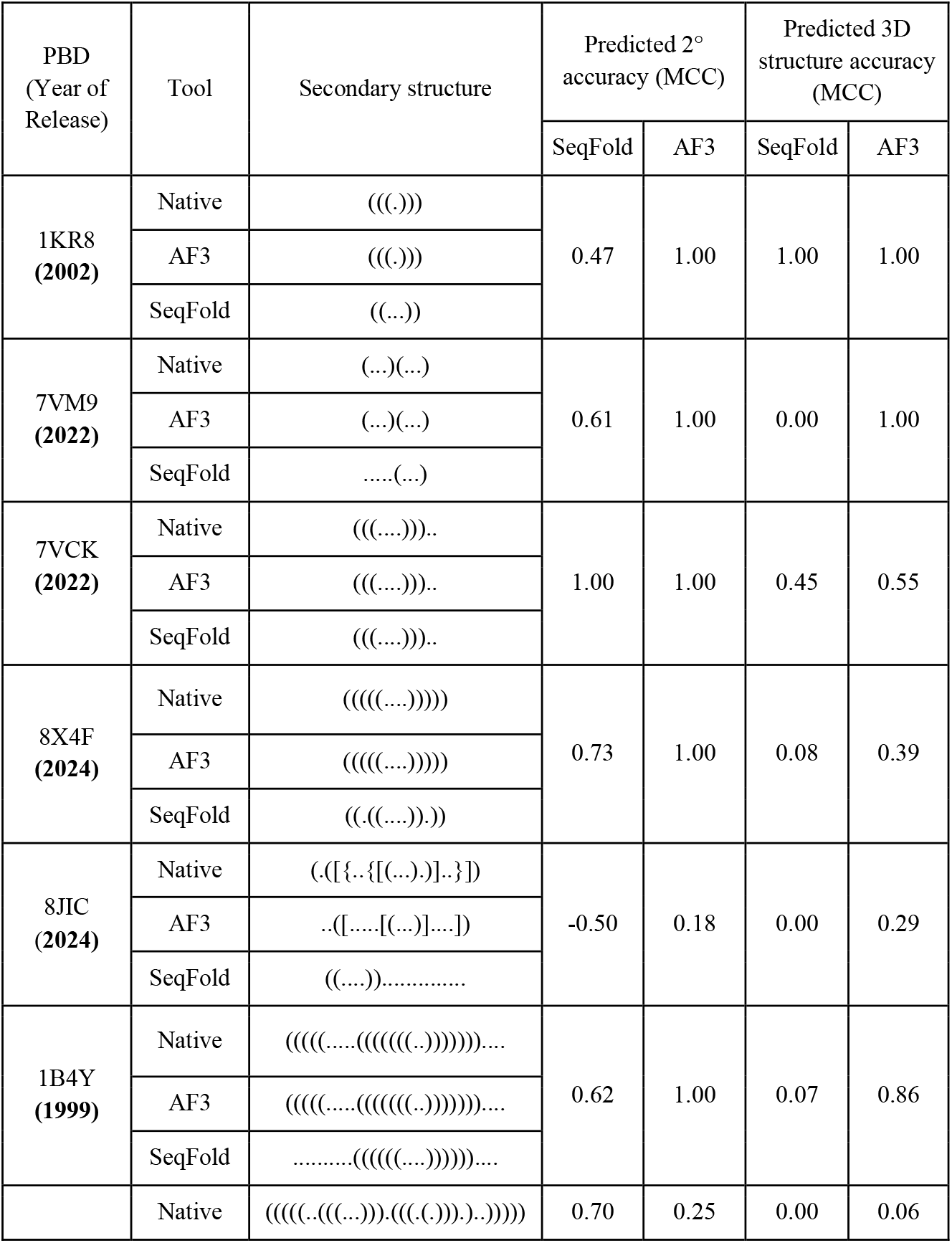
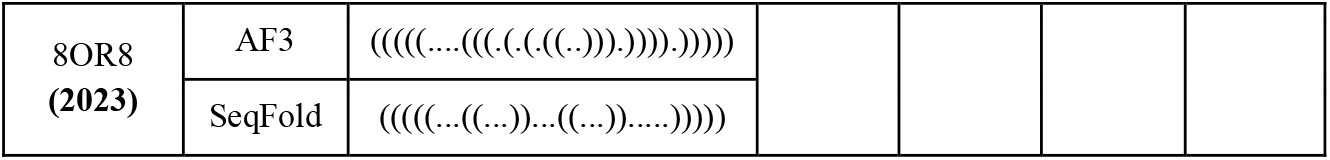
Comparison of SeqFold and AF3 regarding ssDNA secondary structure predictions. 14% (1 out of 7 MD improved cases) and 28% (10 out of 36-case full dataset) secondary structures of SeqFold predictions have the correct secondary structure (MCC = 1.00), compared to 71% (5 out of 7 MD improved cases) and 78% (28 out of 36) cases predicted correctly using AF3. 14% (1 out of 7) and 8% (3 out of 36) of the tertiary structure was predicted correctly using Fold/MMB (3Å cutoff GDT = 1.00), compared to 29% (2 out of 7) and 56% (20 out of 36) cases predicted correctly using AF3. Refer to Table S2 for detailed scores of the full 36 cases.

A correlation was observed between the accuracy of the AF3 predicted secondary structure and the tertiary structure GDT score **(Figure 4)**. The ssDNAs were divided into two categories: those in the AF3 training regime and the ones outside of the training regime. The results showed that an accurate AF3 predicted structure with a high GDT score would have an accurate secondary structure. Examples include 20 out of 36 entries examined **(Table S3)**, ranging from 7 to 27 nucleotides long, had GDT scores of 1.00 and correct secondary structure. When using AF3 to predict the PDB entries that were published after September 2021, AF3 showed potential for predicting fairly accurate secondary structures, but the GDT scores of the overall predicted structures decreased with the increase in length of the ssDNAs **(Figure 4)**.

**Figure 4.**
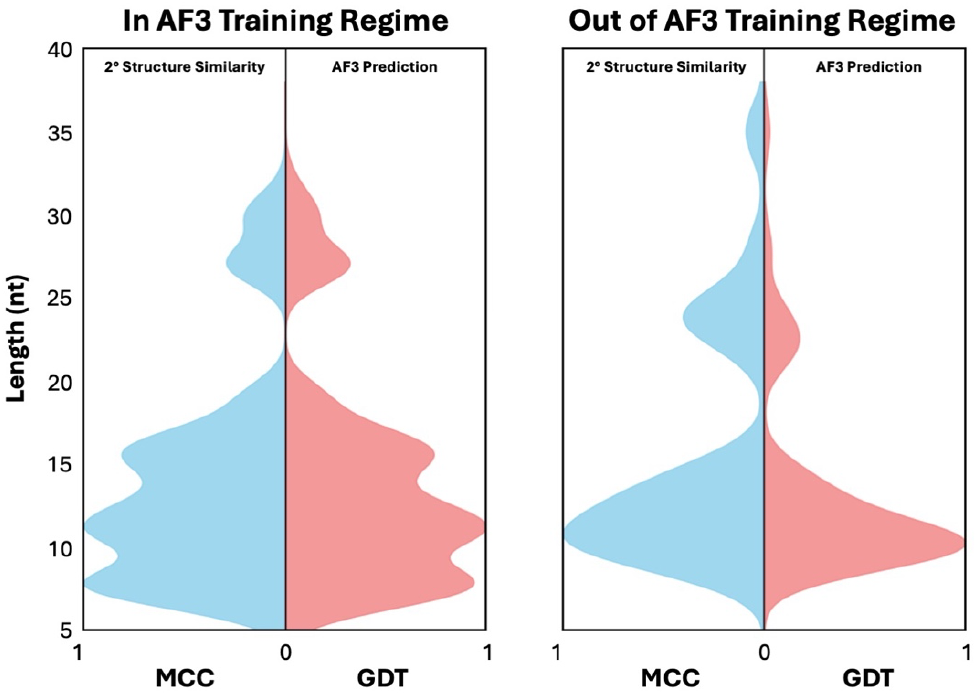
The AF3 predictions (GDT) and their secondary structure accuracy (MCC) of the aptamers in and out of the training regime. nt: nucleotide. The data is smoothed using kernel density estimation (KDE).

Overall, AF3 provides good secondary structure prediction for ssDNA structures. The generated tertiary structures should be used with caution, especially with longer ssDNA cases.

### AF3 predicted structure accuracies of different structure motifs

Simple hairpin predictions showed great accuracy with structures that were a part of its training regime; 14 hairpin structures were predicted with a 1.00 GDT score, and the structures that were in the training regime all had GDT scores above 0.6 **(Table S3)**. The ssDNA with repeats were both part of the AF3 training regime, and they showed high accuracy – 5GWL, a CCTG repeat, and 5GWQ, a TTTA repeat, had scores of 1.00 and 0.86, respectively^33^. The same can be observed with G triplex predictions, in which the two cases, 2MKM and 2MKO, were both in the AF3 training regime, yielding scores of 1.00 and 0.90, respectively.^40,41^ Minidumbbell outside of the AF3 training regime showed equal accuracies as the ones in the training set. The triple helix predicted was also in the AF3 training regime and had a GDT score above 0.8, showing decent accuracy. AF3 predicted G4 results had the widest distribution, from 0.17 to 1.0. The G4 in its training regime was predicted (7E5P) correctly, but AF3 failed to predict the cases outside of its training set: 8JIC and 8RZX had scores of 0.28 and 0.17, respectively.^27,51^ Despite the low GDT scores, the AF3 predicted secondary structure of 8RZX was correct with a MCC score of 1. The low GDT was due to inaccuracies in its flexible bulge region. The error in 8JIC was mainly contributed by the direction of the stacking of the guanines; the AF3-predicted 8JIC was an anti-parallel chair G4, but the correct structure was an anti-parallel basket G4.^60^ AF3 successfully identified them as G4s but failed to provide the exact tertiary structure. The junctions were predicted as hairpins and had the smallest distributions, in which both cases were not a part of the AF3 training regime, and the scores of the predictions were 0.074 and 0.059 for 7QB3 and 8OR8, respectively.^28,54^ AF3 was trained with large quantity of dsDNA helices, in the case of junction predictions, the predicted structures resemble such structure and were simple ssDNA hairpins **(Figure 5, Table S3)**.

**Figure 5.**
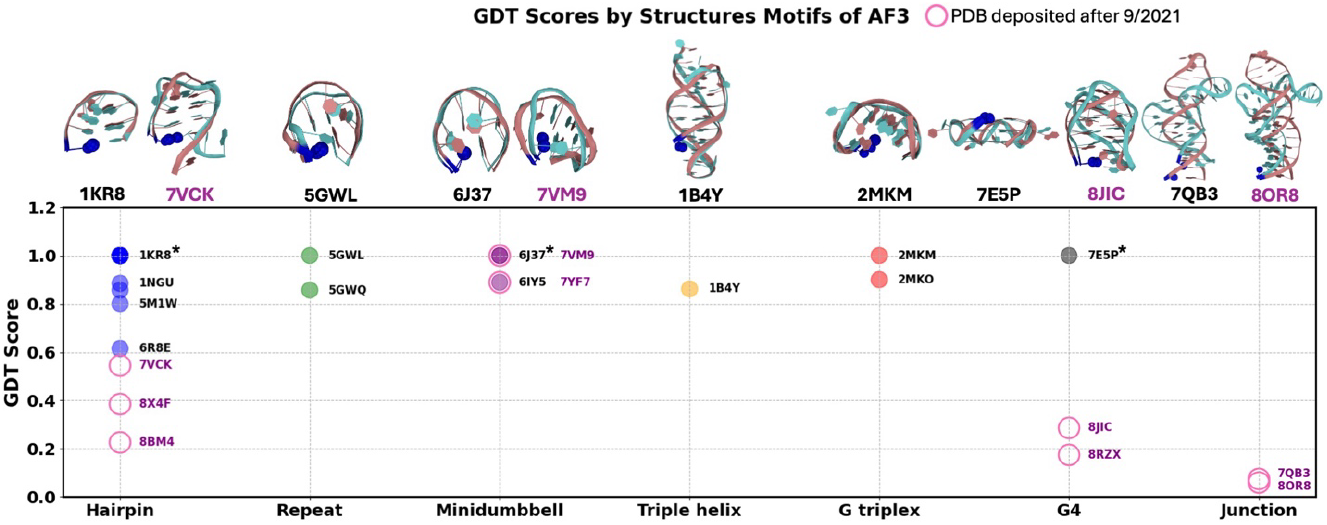
GDT scores of different structure motifs. Pink circles: PDB entries after September 2021, the AF3 training dataset cutoff. Hairpin: Blue. Repeat: Green, 5GWL (CCTG repeats) and 5GWQ (TTTA repeats). Minidumbbell: Plum. Triple helix: Yellow. G triplex: Red. G quadruplex (G4): Black. Junction: Gray. * Indicates multiple entries scored the same and overlapping in the figure.

Overall, AF3 performed well in predicting structures in its training regime; however, the accuracy for ssDNA structures outside of this regime was significantly reduced. AF3 was able to identify hairpin and G4 when they are present and had decent accuracy when predicting their secondary structures. It is a good tool for predicting short minidumbbell motifs, but in this current state, it has limitations when identifying and predicting ssDNA junctions **(Figure 5, Table S3)**.

### AF3 predicted structures can be improved by MD simulations

Two force fields, AMBER99 and OPLS5, showed similar accuracy in our comparison. 1 ns minimization showed that both force fields were reliable, as the structure did not unfold. Overall, the two force fields have very similar performances **(Table 2)**. OPLS5 was selected for further analysis.

**Table 2.** GDT scores of MD optimized AF3 predicted structures with AMBER99 and OPLS5 force fields. The simulation times were 1, 10, and 100 ns. AMBER99 and OPLS5 results are comparable. After 5 10 ns simulations, 14%, 29%, and 43% cases were predicted correctly using Fold/MMB/MD, AF3 + AMBER99sb-idln, and AF3 + OPLS5, respectively. 14%, 29%, and 29% cases were predicted correctly after 100 ns MD using the strategies mentioned prior respectively.

| PBD<br>(Year) | Base<br># | AF3 | Fold+MMB/<br>MD (AMBER99sb-idln,<br>1ns/10ns/100ns) | AF3 + MD<br>(AMBER99sb-idln,<br>1ns/10ns/100ns) | AF3 + MD (OPLS5, 1ns/10<br>ns/100 ns) |
| --- | --- | --- | --- | --- | --- |
| 1KR8<br>(2002) | 7 | 1.00 | 1.00 / 1.00 / 1.00 | 1.00 / 1.00 / 1.00 | 1.00 / 1.00 / 1.00 |
| 7VM9<br>(2021) | 10 | 1.00 | 0.00 / 0.22 / 0.56 | 1.00 / 1.00 / 1.00 | 1.00 / 1.00 / 1.00 |
| 7VCK<br>(2021) | 12 | 0.55 | 0.45 / 0.91 / 0.64 | 0.63 / 0.63 / 0.82 | 0.82 / 0.91 / 0.91 |
| 8X4F<br>(2023) | 14 | 0.39 | 0.08 / 0.15 / 0.31 | 0.61 / 0.92 / 0.92 | 0.85 / 1.00 / 0.92 |
| 8JIC<br>(2024) | 27 | 0.29 | 0.00 / 0.05 / 0.14 | 0.29 / 0.34 / 0.38 | 0.38 / 0.38 / 0.38 |
| 1B4Y<br>(1999) | 30 | 0.86 | 0.00/ 0.00 / 0.17 | 0.87 / 0.87 / 0.90 | 0.90 / 0.93 / 0.93 |
| 8OR8<br>(2023) | 35 | 0.06 | 0.00 / 0.09 / 0.12 | 0.06 / 0.09 / 0.09 | 0.06 / 0.15 / 0.12 |

Three MD strategies were examined using the OPLS5 force field: 10 ns of only the first AF3 prediction (100,000 frames), 10 ns of all five AF3 predictions (100,000 frames/run, 500,000 frames in total), and 100 ns of the first AF3 prediction (500,000 frames). The results showed that running 10 ns of each of the AF3 outputs gives similar results as running 100 ns of only the top AF3 result and is much better than 10 ns of only the top AF3 results **(Figure S1)**. Additionally, the GDT scores of 10 ns of all AF3 outputs and 100 ns of the top output for 7VCK, 8X4F, 8JIC, 1B4Y, and 8OR8 were 0.91/0.91, 1.00/0.92, 0.38/0.38, 0.93/0.93, and 0.15/0.12, respectively **(Figure 6)**. In conclusion, the comparison between 10 and 100 ns MD sampling of only the top AF3 results, and 10 ns of all AF3 results — totaling 50 ns from all AF3 outputs — showed that 10 ns of all AF3 outputs is comparable to running a single 100 ns of the top AF3. Additionally, AMBER99 and OPLS5 shared similar accuracies **(Table 2, Figure S1)**.

**Figure 6.**
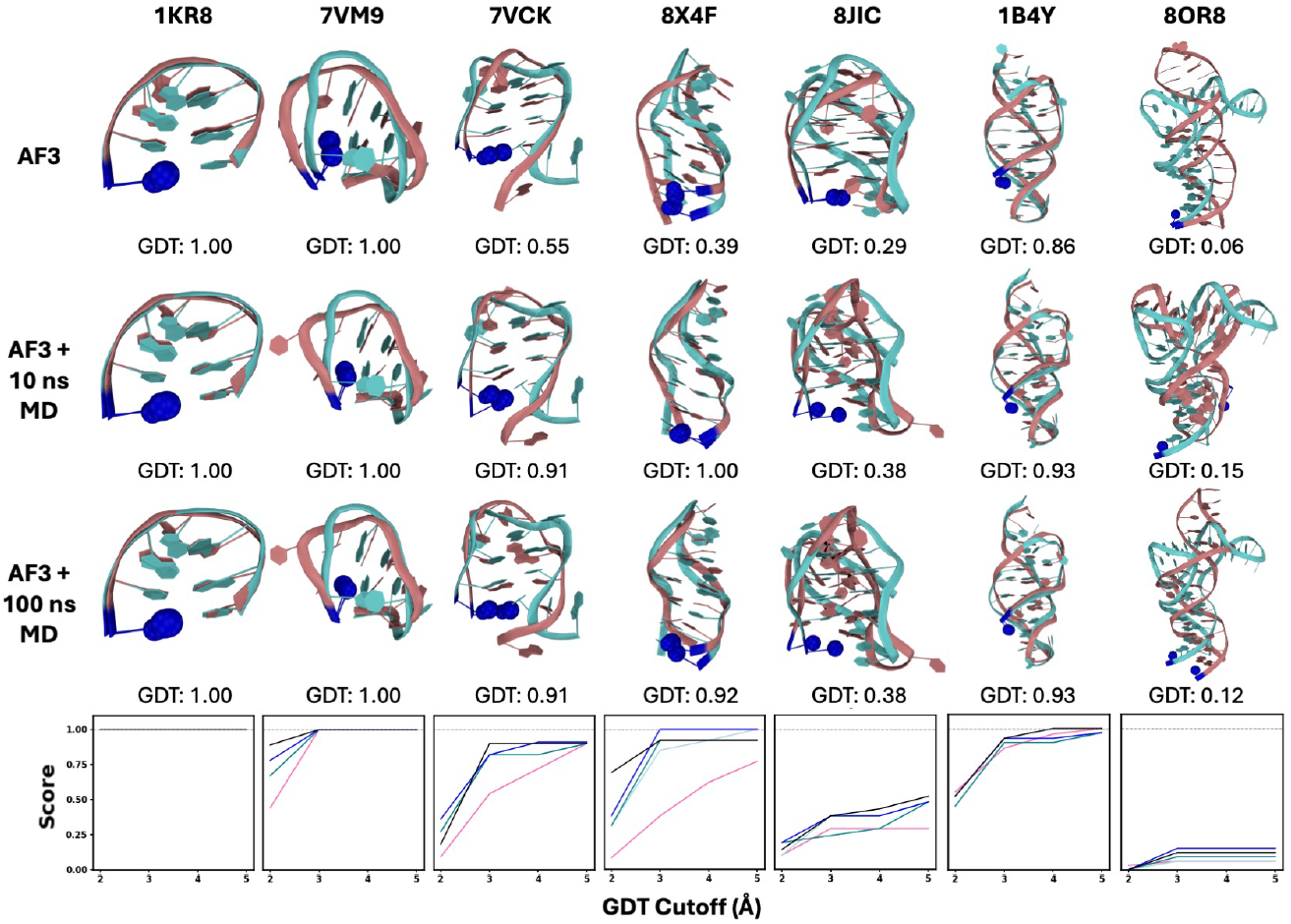
AF3 structure predictions with and without MD simulations in comparison to native aptamer structures. Blue spheres indicate the 5’ end. Line chart: the change of GDT scores with the increase of GDT cutoff. Pink: top AF3 results. Light blue: 1 ns minimization MD of the top AF3 results using OPLS5. Teal: 10 ns MD of the top AF3 results using OPLS5. Blue: 10 ns MD of all the AF3 results using OPLS5. Black: 100 ns MD of the top AF3 results using OPLS5.

Both strategies, Fold/MMB and AF3, were improved by MD simulations. Fold/MMB predicted 7VM9 and 7VCK showed improvements after MD optimizations. AF3 predictions of 7VCK and 8X4F were also greatly improved after 100 ns of MD simulations **(Figure 6)**. The AF3 prediction of 8X4F was very similar to the native structure initially, and all the base pairings were predicted correctly. The MD simulations sampled more backbone conformations, thereby drastically improving the GDT scores from 0.39 to 1.00. Similarly, due to the flexibility of the backbone and the large sample size produced by MD, the GDT score of 7VCK improved from 0.55 to 0.91 **(Figure 6, Table 1)**. The native structure of 8JIC is an antiparallel backet quadruplex (G4), but the predicted structure is an antiparallel chair G4. Improvements with MD was minimal in this case **(Figure 6, Table S2)**. The AF3 result of 8OR8 with the highest GDT score did not have a junction motif rather its prediction was a hairpin. In this case, the 100 ns MD of the best AF3 output sampled only different conformations of the hairpin motif. However, 2 out of the 5 AF3 outputs which did not score the highest were junctions, and they were sampled using the 10 ns MD of all 5 AF3 outputs strategy; the conformation with the highest GDT score using this approach was a junction similar to the native structure **(Figure 6)**. Although its GDT score is low, the junction motif was successfully identified from the poll. Based on the observation, the performance of MD is limited with AF3 predictions with incorrect nucleotide pairing. The correct pairing information or secondary structure is essential for predicting DNA aptamer structures.

### Improving 8X4F AF3 structure with MD simulations

The AF3 predicted 8X4F structure was greatly improved with MD simulations. The 5 AF3 outputs share very similar GDT scores, and the best score was 0.39 **(Figure 7. step 2)**. After 5 10 ns MD simulations, the score improved to 1.00. The MD sampled 500,000 different backbone conformations, and the best conformation for both the 10 ns and 100 ns simulations were almost identical to the native backbone conformation **(Figure 7. step 4)**. The all-atom RMSD of the trajectory was also calculated for all the MDs. The all-atom RMSD showed inverse correlation with the GDT score. Therefore, GDT and RMSD could both be valid benchmarking parameters for ssDNA structure analysis.

**Figure 7.**
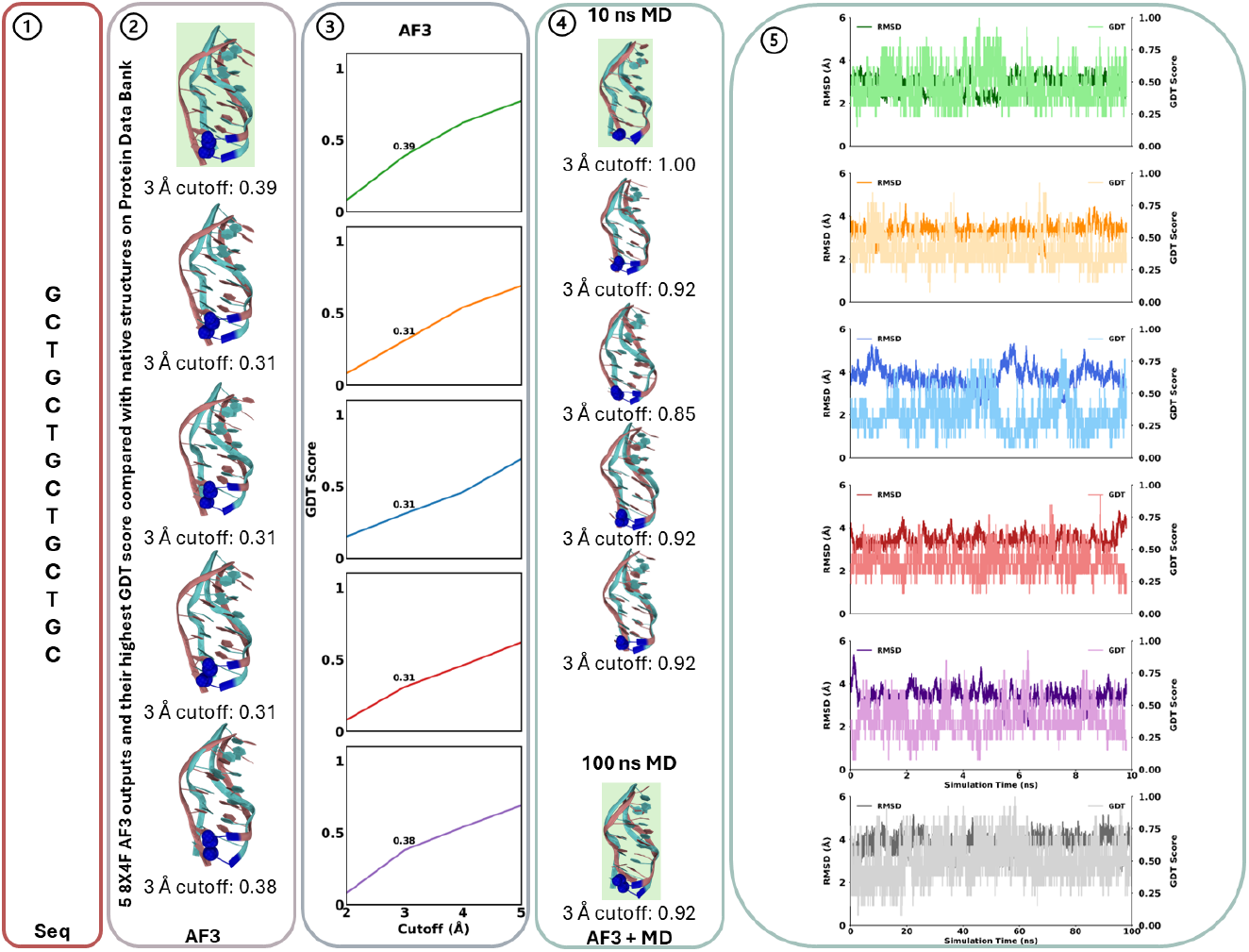
2) Five 8X4F AF3 outputs and their highest GDT score compared to the native structures deposited to PDB. The highest GDT score structure is highlighted in green. 3) The increase of the GDT scores with the increase in cutoffs. 4) the highest GDT score from10 ns of MD of each of the AF3 output and 100 ns of MD of the best AF3 output (background color: light green). 5) RMSD (darker line) and GDT (lighter line) calculated against the native structure.

Both MD strategies — short simulation of multiple AF3 outputs (50 ns sampling) and long simulation of a single AF output (100 ns sampling) — seem to provide sufficient sampling for ssDNA backbone refinement. Given the total sampling strength, 10 ns of all AF3 outputs (in this work, 5 structures) appears to be a more efficient approach.

## Discussion

### Traditional approaches show limited accuracy in both steps

Our data showed that the secondary structure predicted by SeqFold was less accurate than the secondary structures predicted by AF3. MMB was a useful tool to construct 3D models with SeqFold predicted paired bases, however it lacks folding information, and the 3D models were hard to optimize with MD. There are cases where SeqFold was more accurate than AF3 and was able to identify the sequence as a junction where AF3 failed. In cases of complex motifs, such as junctions, SeqFold secondary structure predictions outperformed AF3 for 7QB3 and 8OR8 **(Figure 8, Table S2)**. Additionally, SeqFold predicted the secondary structure of long hairpin 8BM4 was more accurate than AF3. However, SeqFold was unable to predict any pairs for 5 sequences: 2MKM (G triplex), 2MKO (G triplex), 5M1W (G hairpin), 6R8E (G hairpin), 1RDE (G4). SeqFold also showed poor accuracy when predicting G4 secondary structures – 7E5P, 8JIC, and 8RZX – their MCC scores were – 0.65, – 0.50, and – 0.35, respectively **(Figure 8, Table S2)**. Interestingly, SeqFold is a decent candidate for junction prediction, but it has poor performance when predicting G-rich structures. In addition to secondary structure prediction accuracy, the 3D structures generated by MMB were also not as accurate as AF3 predictions **(Table 1)**.

**Figure 8.**
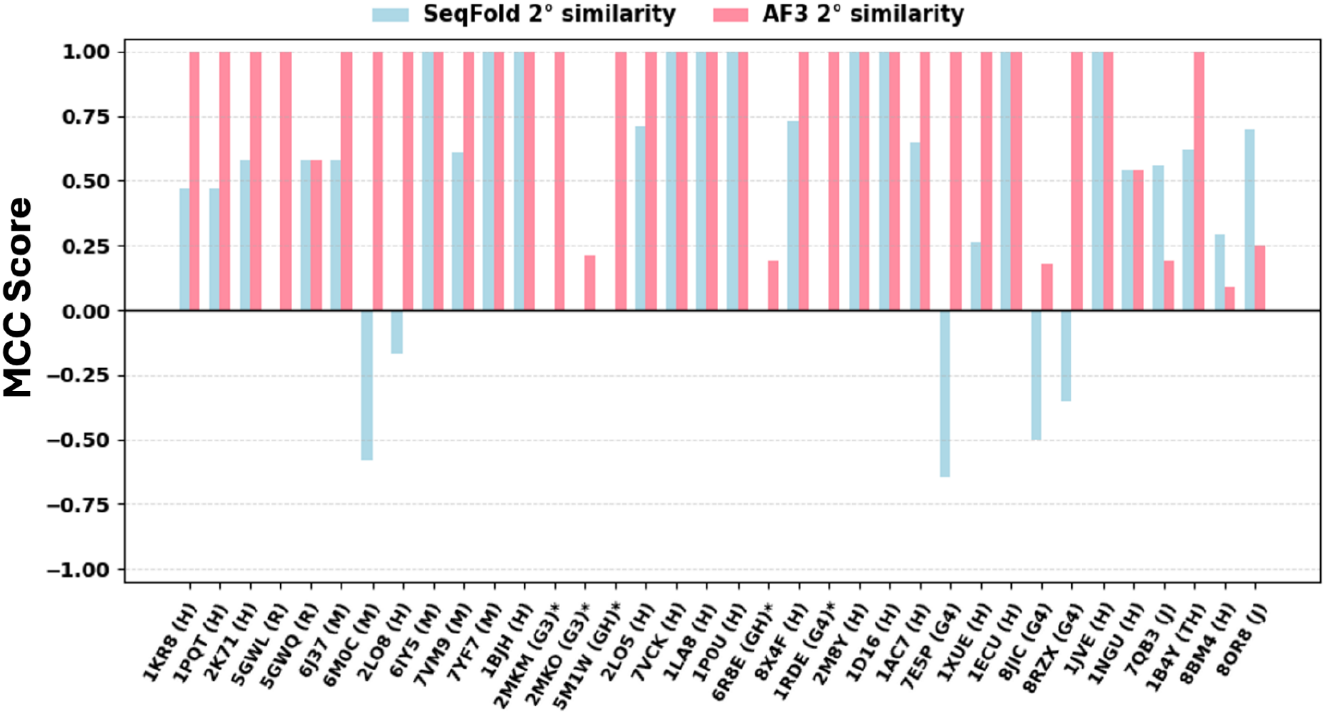
MCC score of the secondary structures predicted by SeqFold (light blue) and AF3 (pink). Structure motifs are indicated with the PDB ID. H: hairpin. R: repeats. M: minidumbbell. G3: G triplex. GH: G hairpin. G4: G-quadruplex. J: junction. TH: triple helix. * Indicates no prediction generated from SeqFold.

### The AF3/MD pipeline is an efficient approach for predicting ssDNA structure

The traditional approach essentially assumes that base pairing of a ssDNA can be predicted without the knowledge of the 3D structure. It is not surprising that its accuracy is dependent on the secondary structure prediction tool. Our data clearly indicates that inaccurate prediction of the secondary structure led to the failure of ssDNA structure prediction, for example, in the case of 8JIC and 8OR8. Furthermore, our data show that AF3 was fairly accurate for secondary structure prediction; its accuracy surpassed the secondary structure prediction tool, SeqFold, with a success rate (MCC = 1.00) of 78% (28 out of 36 predicted correctly) and 28% (10 out of 36 predicted correctly), respectively **(Table 1, Table S2)**.

However, we also found limitations in AF3’s ability to detect complex DNA motifs, including G4 and junctions. For example, 8JIC and 8RXZ are found to adopt the G4 structures. 8JIC is an anti-parallel basket G4, but the predicted structure is an anti-parallel chair G4 **(Figure 6)**. This difference led to a mismatch in the base pairings and the predicted base pairs were 1 nucleotide off from the native pairings, in which resulted in a low MMC score **(Table 1)**. Despite the low MCC and GDT scores, AF3 was able to identify 8JIC as a G4 **(Figure 5)**. 8RXZ was predicted very well with 100% correct base pairing predictions. The native and predicted structures were both parallel G4s **(Table S3)**, and the errors in the predicted 8RXZ originated from its unpaired flexible loop. Overall, AF3 can effectively identify G4 when it is present, and it is an efficient tool for G4 secondary structure predictions. DNA junctions are structures where two or more DNA duplex stems connect, such as the three-way and four-way junctions found in DNAzymes to control their catalytic activity. For junction structures like 7QB3 and 8OR8, they were often predicted as simple hairpins by AF3 **(Table 1, Figure 5)**. The AF3 training regime likely lacks ssDNA junction structures, resulting in the failure to identify junctions or provide accurate secondary structures.

Once AF3 predicts the correct secondary structure, MD simulations can efficiently sample many relevant backbone conformations. Our best example to illustrate the power of MD to improve AF3 prediction is the case of 8X4F, for which the GDT score improved from 0.38 to 1.00 after MD sampling **(Figure 7)**. For large improvements like this, AF3 is required to provide a correct output with all the right base pairings. However, an inverse relationship was observed between the length of the ssDNA and the prediction accuracy, with the secondary and tertiary structure prediction of longer sequences being less accurate **(Figure 3)**.

### AF3 may be biased toward common B-form DNA

As discussed above, the long and complex ssDNA, such as junctions, could be predicted as long hairpins by AF3, for example, 8OR8 and 7QB3. We are aware that the AF3-predicted hairpin structures, such as 8BM4 and 1NGU, resemble B-form DNAs, and the angle distributions of the AF3-predicted hairpin structures are similar to those of B-form DNAs, presumably caused by the bias in the AF3 training regime. As shown in Figure 8, the dihedral angles from the four cases previously mentioned were plotted and compared to those of A, B, and Z-form DNA (analysis of a manually curated dataset from PDB in summer 2022). The dihedral angle distribution of the AF3-predicted hairpins most closely resembled those of the B-form DNA, while the alpha, delta, and zeta angles clearly differ between the AF3-predicted structure and A and Z-form DNAs. As ssDNA hairpins consist of a B-form stem and a short loop, it is not difficult to understand why AF3 can be accurate in predicting ssDNA hairpin structures **(Figure 9, Table S2)**. However, this bias reduces confidence in predicting other ssDNA motifs, particularly junction motifs.

**Figure 9.**
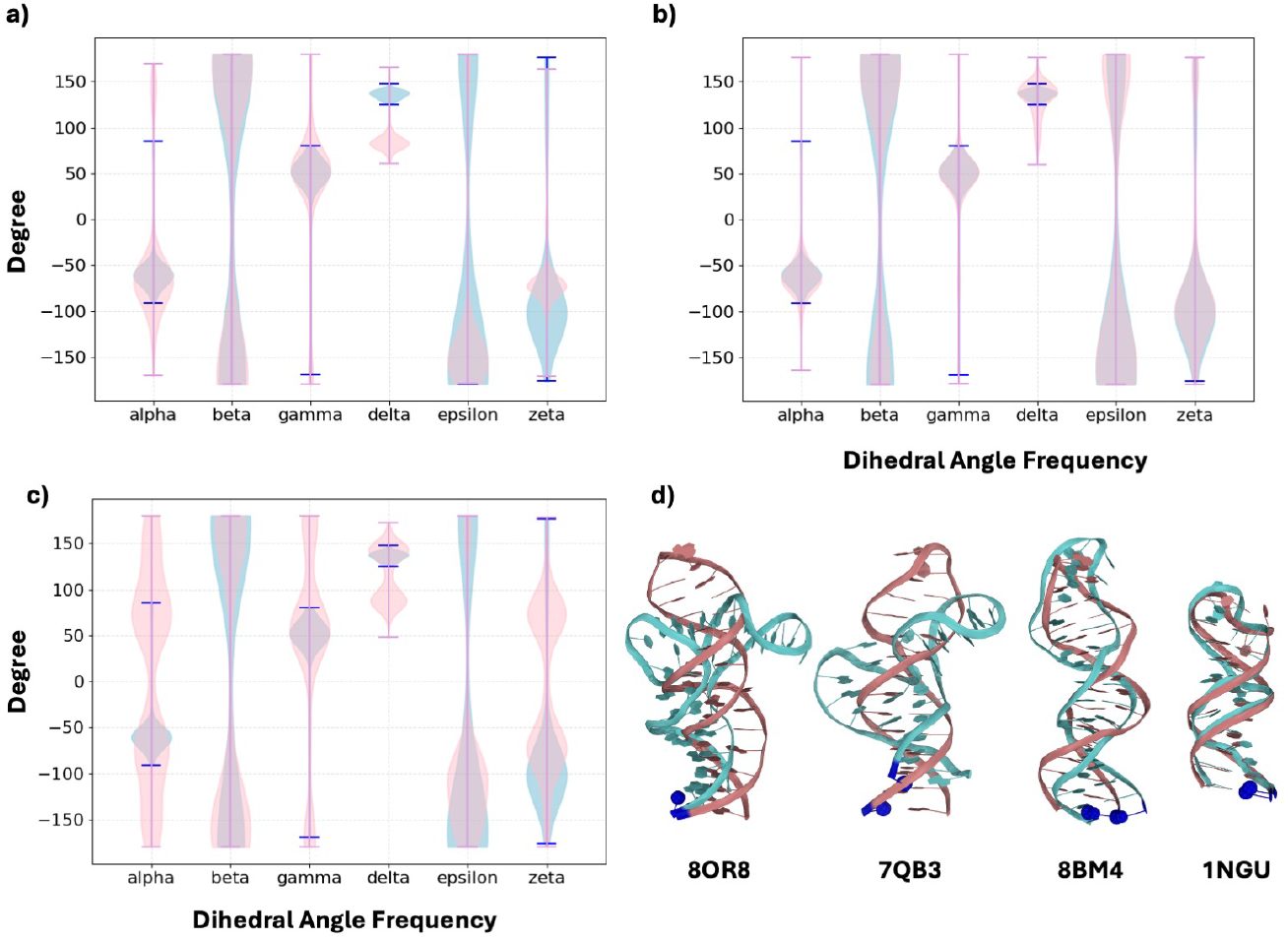
Violin plot of AF3 predicted structure dihedral angle distribution (light blue) and different forms of DNA dihedral angle distribution (pink, analysis of our in-house dataset of 64 A-form, 66 B-form, and 151 Z-form DNAs collected from PDB by June 2022). a) A-form DNA dihedral angle and AF3 predicted structure dihedral angle distributions. b) B-form DNA dihedral angle and AF3 predicted structure dihedral angle distributions. c) Z-form DNA dihedral angle and AF3 predicted structure dihedral angle distributions. d) aligned structures of AF3 predictions with the highest GDT (pink) with the native structure (light blue), blue spheres: the 5’ end of the sequences.

## Conclusion

Small ssDNA structure prediction is critical for the structure-based design of DNA therapeutics. In this work, we have investigated two strategies for DNA structure prediction and carefully evaluated their predictions with multiple metrics. The traditional approach, which employs classical secondary structure prediction and 3D structure conversion, displayed limited accuracy in both steps and thus resulted in a low success rate in our study. We found that AF3 outperformed the traditional approach for predicting both secondary structures and 3D structures; however, its accuracy decreased with an increase in sequence length. Interestingly, our data suggest that all-atom MD simulation with commonly used force fields can improve the predicted structure from either the traditional approach or the AF3 approach. A further analysis of the performance with various ssDNA folding motifs revealed a potential bias of AF3 toward the B-form DNA conformation. Improvement on other types of structure, such as G4 quadruplex and DNA junction, are still needed for future application to ssDNA therapeutic design.

## Acknowledgments

We thank Prof. Tom Soh and Hajime Fujita (Stanford University) and Prof. Hanadi Sleiman (McGill University) for helpful discussions. This work was supported by an NSF CAREER award (CHE-2410514 to J.L.) and the AnalytiXIN fellowship.

## Data Availability

The data and GDT analysis scripts are available at https://github.com/jianingli-purdue/ssDNA-scoring.

## Reference

1. Rinaldi, C.; Wood, M. J. A. Antisense Oligonucleotides: The next Frontier for Treatment of Neurological Disorders. Nat Rev Neurol 2018, 14 (1), 9–21.

2. Egli, M.; Manoharan, M. Chemistry, Structure and Function of Approved Oligonucleotide Therapeutics. Nucleic Acids Res 2023, 51 (6), 2529–2573.

3. Vinjamuri, B. P.; Pan, J.; Peng, P. A Review on Commercial Oligonucleotide Drug Products. J Pharm Sci 2024, 113 (7), 1749–1768.

4. Roberts, T. C.; Langer, R.; Wood, M. J. A. Advances in Oligonucleotide Drug Delivery. Nat Rev Drug Discov 2020, 19 (10), 673–694.

5. Zhou, J.; Rossi, J. Aptamers as Targeted Therapeutics: Current Potential and Challenges. Nature reviews. Drug discovery 2016, 16 (3), 181.

6. Huang, Z.; Niu, L. RNA Aptamers for AMPA Receptors. Neuropharmacology 2021, 199, 108761.

7. He, A.; Wan, L.; Zhang, Y.; Yan, Z.; Guo, P.; Han, D.; Tan, W. Structure-Based Investigation of a DNA Aptamer Targeting PTK7 Reveals an Intricate 3D Fold Guiding Functional Optimization. Proceedings of the National Academy of Sciences 2024, 121 (29), e2404060121.

8. Lyu, M.; Chan, C. H.; Chen, Z.; Liu, Y.; Yu, Y. Advantages, Applications, and Future Directions of in Vivo Aptamer SELEX: A Review. Molecular Therapy. Nucleic Acids 2025, 36 (3), 102575.

9. Odeh, F.; Nsairat, H.; Alshaer, W.; Ismail, M. A.; Esawi, E.; Qaqish, B.; Bawab, A. A.; Ismail, S. I. Aptamers Chemistry: Chemical Modifications and Conjugation Strategies. Molecules 2019, 25 (1), 3.

10. Cui, Y.; Chen, Y.; Zhang, Y.; Zhang, L. Advances in Aptamer Technology for Target-Based Drug Discovery. Journal of Pharmaceutical Analysis 2025, 101369.

11. Bonilla, S. L.; Jang, K. Challenges, Advances, and Opportunities in RNA Structural Biology by Cryo-EM. Current Opinion in Structural Biology 2024, 88, 102894.

12. Singh, J.; Hanson, J.; Paliwal, K.; Zhou, Y. RNA Secondary Structure Prediction Using an Ensemble of Two-Dimensional Deep Neural Networks and Transfer Learning. Nat Commun 2019, 10 (1), 5407.

13. Shen, T.; Hu, Z.; Sun, S.; Liu, D.; Wong, F.; Wang, J.; Chen, J.; Wang, Y.; Hong, L.; Xiao, J.; Zheng, L.; Krishnamoorthi, T.; King, I.; Wang, S.; Yin, P.; Collins, J. J.; Li, Y. Accurate RNA 3D Structure Prediction Using a Language Model-Based Deep Learning Approach. Nat Methods 2024, 21 (12), 2287–2298.

14. Kamga Youmbi, F. I.; Kengne Tchendji, V.; Tayou Djamegni, C. P-FARFAR2: A Multithreaded Greedy Approach to Sampling Low-Energy RNA Structures in Rosetta FARFAR2. Computational Biology and Chemistry 2023, 104, 107878.

15. Abramson, J.; Adler, J.; Dunger, J.; Evans, R.; Green, T.; Pritzel, A.; Ronneberger, O.; Willmore, L.; Ballard, A. J.; Bambrick, J.; Bodenstein, S. W.; Evans, D. A.; Hung, C.-C.; O’Neill, M.; Reiman, D.; Tunyasuvunakool, K.; Wu, Z.; Žemgulytė, A.; Arvaniti, E.; Beattie, C.; Bertolli, O.; Bridgland, A.; Cherepanov, A.; Congreve, M.; Cowen-Rivers, A. I.; Cowie, A.; Figurnov, M.; Fuchs, F. B.; Gladman, H.; Jain, R.; Khan, Y. A.; Low, C.M. R.; Perlin, K.; Potapenko, A.; Savy, P.; Singh, S.; Stecula, A.; Thillaisundaram, A.; Tong, C.; Yakneen, S.; Zhong, E. D.; Zielinski, M.; Žídek, A.; Bapst, V.; Kohli, P.; Jaderberg, M.; Hassabis, D.; Jumper, J. M. Accurate Structure Prediction of Biomolecular Interactions with AlphaFold 3. Nature 2024, 630 (8016), 493–500.

16. Kilgour, M.; Liu, T.; Walker, B. D.; Ren, P.; Simine, L. E2EDNA: Simulation Protocol for DNA Aptamers with Ligands. J. Chem. Inf. Model. 2021, 61 (9), 4139–4144.

17. Cortes, M.; Sun, X.; Anusha; Joseph Batchelder-Schwab, E.; Li, J.; Siraj, N.; Jampana, R.; Zhang, Y.; Bai, Y.; Mao, C. AlphaFold 3 Modeling of DNA Nanomotifs: Is It Reliable? Nanoscale Horizons 2025, 10 (7), 1428–1435.

18. Bergonzo, C.; Grishaev, A. Critical Assessment of RNA and DNA Structure Predictions via Artificial Intelligence: The Imitation Game. J. Chem. Inf. Model. 2025, 65 (7), 3544–3554.

19. Ouyang, Z.; Snyder, M. P.; Chang, H. Y. SeqFold: Genome-Scale Reconstruction of RNA Secondary Structure Integrating High-Throughput Sequencing Data. Genome Res 2013, 23 (2), 377–387.

20. Flores, S. C.; Sherman, M. A.; Bruns, C. M.; Eastman, P.; Altman, R. B. Fast Flexible Modeling of RNA Structure Using Internal Coordinates. IEEE/ACM Trans Comput Biol Bioinform 2011, 8 (5), 1247–1257.

21. Li, W.; Schaeffer, R. D.; Otwinowski, Z.; Grishin, N. V. Estimation of Uncertainties in the Global Distance Test (GDT_TS) for CASP Models. PLoS ONE 2016, 11 (5), e0154786.

22. Ochoa, S.; Milam, V. T. Direct Modeling of DNA and RNA Aptamers with AlphaFold 3: A Promising Tool for Predicting Aptamer Structures and Aptamer–Target Interactions. ACS Synth. Biol. 2025.

23. Padrta, P.; Štefl, R.; Králík, L.; Žídek, L.; Sklenář, V. Refinement of d(GCGAAGC) Hairpin Structure Using One- and Two-Bond Residual Dipolar Couplings. J Biomol NMR 2002, 24 (1), 1–14.

24. Ngai, C. K.; Lam, S. L.; Lee, H. K.; Guo, P. A Purine and a Backbone Discontinuous Site Alter the Structure and Thermal Stability of DNA Minidumbbells Containing Two Pentaloops. FEBS Letters 2022, 596 (6), 826–840.

25. Wan, L.; He, A.; Li, J.; Guo, P.; Han, D. High-Resolution NMR Structures of Intrastrand Hairpins Formed by CTG Trinucleotide Repeats. ACS Chem. Neurosci. 2024, 15 (4), 868–876.

26. van Dongen, M. J. P.; Doreleijers, J. F.; van der Marel, G. A.; van Boom, J. H.; Hilbers, C. W.; Wijmenga, S. S. Structure and Mechanism of Formation of the H-Y5 Isomer of an Intramolecular DNA Triple Helix. Nat Struct Mol Biol 1999, 6 (9), 854–859.

27. Galer, P.; Wang, B.; Plavec, J.; Šket, P. Unveiling the Structural Mechanism of a G-Quadruplex pH–Driven Switch. Biochimie 2023, 214, 73–82.

28. Wieruszewska, J.; Pawłowicz, A.; Połomska, E.; Pasternak, K.; Gdaniec, Z.; Andrałojć, W. The 8-17 DNAzyme Can Operate in a Single Active Structure Regardless of Metal Ion Cofactor. Nat Commun 2024, 15 (1), 4218.

29. Yi, J.; Wan, L.; Liu, Y.; Lam, S. L.; Chan, H. Y. E.; Han, D.; Guo, P. NMR Solution Structures of d(GGCCTG)n Repeats Associated with Spinocerebellar Ataxia Type 36. Int J Biol Macromol 2022, 201, 607–615.

30. Chattopadhyaya, R.; Grzeskowiak, K.; Dickerson, R. E. Structure of a T4 Hairpin Loop on a Z-DNA Stem and Comparison with A-RNA and B-DNA Loops. Journal of Molecular Biology 1990, 211 (1), 189–210.

31. Padrta, P.; Štefl, R.; Králík, L.; Žídek, L.; Sklenář, V. Refinement of d(GCGAAGC) Hairpin Structure Using One- and Two-Bond Residual Dipolar Couplings. J Biomol NMR 2002, 24 (1), 1–14.

32. Santini, G. P. H.; Cognet, J. A. H.; Xu, D.; Singarapu, K. K.; Hervé du Penhoat, C. Nucleic Acid Folding Determined by Mesoscale Modeling and NMR Spectroscopy: Solution Structure of d(GCGAAAGC). J. Phys. Chem. B 2009, 113 (19), 6881–6893.

33. Guo, P.; Lam, S. L. Minidumbbell: A New Form of Native DNA Structure. J. Am. Chem. Soc. 2016, 138 (38), 12534–12540.

34. Unprecedented hydrophobic stabilizations from a reverse wobble T·T mispair in DNA minidumbbell: Journal of Biomolecular Structure and Dynamics: Vol 38, No 7 - Get Access.

35. Ngai, C. K.; Lam, S. L.; Lee, H. K.; Guo, P. High-Resolution Structures of DNA Minidumbbells Comprising Type II Tetraloops with a Purine Minor Groove Residue. J. Phys. Chem. B 2020, 124 (25), 5131–5138.

36. del Mundo, I. M. A.; Siters, K. E.; Fountain, M. A.; Morrow, J. R. Structural Basis for Bifunctional Zinc(II) Macrocyclic Complex Recognition of Thymine Bulges in DNA. Inorg. Chem. 2012, 51 (9), 5444–5457.

37. Guo, P.; Lam, S. L. Minidumbbell Structures Formed by ATTCT Pentanucleotide Repeats in Spinocerebellar Ataxia Type 10. Nucleic Acids Research 2020, 48 (13), 7557–7568.

38. Li, J.; Wan, L.; Wang, Y.; Chen, Y.; Lee, H. K.; Lam, S. L.; Guo, P. Solution Nuclear Magnetic Resonance Structures of ATTTT and ATTTC Pentanucleotide Repeats Associated with SCA37 and FAMEs. ACS Chem. Neurosci. 2023, 14 (2), 289–299.

39. Chou, S.-H.; Zhu, L.; Gao, Z.; Cheng, J.-W.; Reid, B. R. Hairpin Loops Consisting of Single Adenine Residues Closed by Sheared A·A and G·G Pairs Formed by the DNA Triplets AAA and GAG: Solution Structure of the d(GTACAAAGTAC) Hairpin. Journal of Molecular Biology 1996, 264 (5), 981–1001.

40. Cerofolini, L.; Amato, J.; Giachetti, A.; Limongelli, V.; Novellino, E.; Parrinello, M.; Fragai, M.; Randazzo, A.; Luchinat, C. G-Triplex Structure and Formation Propensity. Nucleic Acids Research 2014, 42 (21), 13393–13404.

41. Gajarský, M.; Živković, M. L.; Stadlbauer, P.; Pagano, B.; Fiala, R.; Amato, J.; Tomáška, L.; Šponer, J.; Plavec, J.; Trantírek, L. Structure of a Stable G-Hairpin. J. Am. Chem. Soc. 2017, 139 (10), 3591–3594.

42. Weisenseel, J. P.; Reddy, G. R.; Marnett, L. J.; Stone, M. P. Structure of the 1,N2-Propanodeoxyguanosine Adduct in a Three-Base DNA Hairpin Loop Derived from a Palindrome in the Salmonella Typhimurium hisD3052 Gene. Chem. Res. Toxicol. 2002, 15 (2), 140–152.

43. Chin, K.-H.; Chou, S.-H. Sheared-Type Ganti·Csyn Base-Pair: A Unique d(GXC) Loop Closure Motif. Journal of Molecular Biology 2003, 329 (2), 351–361.

44. Živković, M. L.; Gajarský, M.; Beková, K.; Stadlbauer, P.; Vicherek, L.; Petrová, M.; Fiala, R.; Rosenberg, I.; Šponer, J.; Plavec, J.; Trantírek, L. Insight into Formation Propensity of Pseudocircular DNA G-Hairpins. Nucleic Acids Research 2021, 49 (4), 2317–2332.

45. Mao, X.; Marky, L. A.; Gmeiner, W. H. NMR Structure of the Thrombin-Binding DNA Aptamer Stabilized by Sr2+. J Biomol Struct Dyn 2004, 22 (1), 25–33.

46. Lim, K. W.; Phan, A. T. Structural Basis of DNA Quadruplex–Duplex Junction Formation. Angewandte Chemie International Edition 2013, 52 (33), 8566–8569.

47. van Dongen, M. J. P.; Mooren, M. M. W.; Willems, E. F. A.; van der Marel, G. A.; van Boom, J. H.; Wijmenga, S. S.; Hilbers, C. W. Structural Features of the DNA Hairpin d(ATCCTA-GTTA-TAGGAT): Formation of a G-A Base Pair in the Loop. Nucleic Acids Research 1997, 25 (8), 1537–1547.

48. Tsukakoshi, K.; Yamagishi, Y.; Kanazashi, M.; Nakama, K.; Oshikawa, D.; Savory, N.; Matsugami, A.; Hayashi, F.; Lee, J.; Saito, T.; Sode, K.; Khunathai, K.; Kuno, H.; Ikebukuro, K. G-Quadruplex-Forming Aptamer Enhances the Peroxidase Activity of Myoglobin against Luminol. Nucleic Acids Research 2021, 49 (11), 6069–6081.

49. Zhu, L.; Chou, S. H.; Reid, B. R. A Single G-to-C Change Causes Human Centromere TGGAA Repeats to Fold Back into Hairpins. Proceedings of the National Academy of Sciences 1996, 93 (22), 12159–12164.

50. Bank, R. P. D. RCSB PDB - 1ECU: SOLUTION STRUCTURE OF E2F BINDING DNA FRAGMENT GCGCGAAAC-T-GTTTCGCGC.

51. Chatterjee, O.; Jana, J.; Panda, S.; Dutta, A.; Sharma, A.; Saurav, S.; Motiani, R. K.; Weisz, K.; Chatterjee, S. Remodeling Ca2+ Dynamics by Targeting a Promising E-Box Containing G-Quadruplex at ORAI1 Promoter in Triple-Negative Breast Cancer. Cell Calcium 2024, 123, 102944.

52. Ulyanov, N. B.; Bauer, W. R.; James, T. L. High-Resolution NMR Structure of an AT-Rich DNA Sequence. J Biomol NMR 2002, 22 (3), 265–280.

53. Shiflett, P. R.; Taylor-McCabe, K. J.; Michalczyk, R.; Silks, L. A. “Pete”; Gupta, G. Structural Studies on the Hairpins at the 3’ Untranslated Region of an Anthrax Toxin Gene,. Biochemistry 2003, 42 (20), 6078–6089.

54. Novotný, A.; Plavec, J.; Kocman, V. Structural Polymorphism Driven by a Register Shift in a CGAG-Rich Region Found in the Promoter of the Neurodevelopmental Regulator AUTS2 Gene. Nucleic Acids Research 2023, 51 (6), 2602–2613.

55. Ferrell, J.; Shi, Y.; Shi, Y.; Schneebeli, S.; Li, J. Tunable Conformational Fluctuations of DNA Nanocages. ChemRxiv January 27, 2025.

56. Markham, N. R.; Zuker, M. UNAFold: Software for Nucleic Acid Folding and Hybridization. Methods Mol Biol 2008, 453, 3–31. 10.1007/978-1-60327-429-6_1.

57. Singh, J.; Hanson, J.; Paliwal, K.; Zhou, Y. RNA Secondary Structure Prediction Using an Ensemble of Two-Dimensional Deep Neural Networks and Transfer Learning. Nat Commun 2019, 10 (1), 5407. 10.1038/s41467-019-13395-9.

58. Fu, L.; Cao, Y.; Wu, J.; Peng, Q.; Nie, Q.; Xie, X. UFold: Fast and Accurate RNA Secondary Structure Prediction with Deep Learning. Nucleic Acids Res 2022, 50 (3), e14. 10.1093/nar/gkab1074.

59. Parisien, M.; Major, F. The MC-Fold and MC-Sym Pipeline Infers RNA Structure from Sequence Data. Nature 2008, 452 (7183), 51–55. 10.1038/nature06684.

60. Meier-Stephenson, V. G4-Quadruplex-Binding Proteins: Review and Insights into Selectivity. Biophys Rev 2022, 14 (3), 635–654. 10.1007/s12551-022-00952-8.

