## Supplemental Information for "Small Single-Stranded DNA Structure Prediction"

### Details of MD Simulation and Analysis:

**Desmond** from Schrodinger, Inc. Software Suites 2024-4 was used for molecular dynamics simulations with **OPLS5** forcefield. Direct AF3 outputs with the highest GDT score compared to the native structure entries from Protein Data Bank were imported as initial structures. SPC water molecules and neutralizing $Mg^{2+}$ ions were constructed. Simulations were performed using the NPT ensemble at 300 K at 1.0 bar. The time step was set at 2 fs, the recording interval was 1 ps, and the number of frames was 100,000.

**GROMACS** was also used for molecular dynamics simulations with **AMBER99sb-idln** forcefield. The same initial structure was used as previously described from the **Desmond** MD setup. SPC water molecules and neutralizing $Mg^{2+}$ were constructed. Simulations were also performed using the NPT ensemble at 300 K at 1.0 bar. The time step was set at 2 fs, the recording interval was 1 ps, and the number of frames was 100,000.

For **Desmond** trajectory analysis, the trajectories were all loaded into **VMD** and converted to gro and trr files and analyzed with MDAnalysis. For **AMBER99** trajectory, the file type conversion was not performed. Each frame of the trajectory from both **Desmond** and **AMBER99** was extracted and was compared to the reference structure – native structure on Protein DataBank. The GDT score of each frame was calculated against the reference structure. The highest GDT score was recorded.

The **AF3** results are combined into a trr trajectory file and each output from AF3 was analyzed as a trr trajectory using the same method stated previously with MD generated trajectories.

**Table S1. Desmond (OPLS5)** and **GROMACS (AMBER99)** MD simulation setup information. SPC water box was generated 10 Å around the solute. Simulation temperature was 300 K.

| PDB ID | Replica Count OPLS5 | Replica Count AMBER99 | Mg2+ Count | Time (ns) |
|---|---|---|---|---|
| 1KR8 **(2002)** | 1 (1 ns)<br>1 (10 ns)<br>2 (100ns) | 1 (1 ns)<br>1 (10 ns)<br>1 (100ns) | 3 | 1,<br>10 of top AF3,<br>10 of all AF3,<br>100 |
| 7VM9 **(2021)** | 1 (1 ns)<br>1 (10 ns)<br>2 (100ns) | 1 (1 ns)<br>1 (10 ns)<br>1 (100ns) | 5 | 1,<br>10 of top AF3,<br>10 of all AF3,<br>100 |
| 7VCK **(2021)** | 1 (1 ns)<br>1 (10 ns)<br>2 (100ns) | 1 (1 ns)<br>1 (10 ns)<br>1 (100ns) | 6 | 1,<br>10 of top AF3,<br>10 of all AF3,<br>100 |
| 8X4F **(2023)** | 1 (1 ns)<br>1 (10 ns)<br>2 (100ns) | 1 (1 ns)<br>1 (10 ns)<br>1 (100ns) | 7 | 1,<br>10 of top AF3,<br>10 of all AF3,<br>100 |
| 8JIC (**2024)** | 1 (1 ns)<br>1 (10 ns)<br>2 (100ns) | 1 (1 ns)<br>1 (10 ns)<br>1 (100ns) | 11 | 1,<br>10 of top AF3,<br>10 of all AF3,<br>100 |
| 1B4Y **(1999)** | 1 (1 ns)<br>1 (10 ns)<br>2 (100ns) | 1 (1 ns)<br>1 (10 ns)<br>1 (100ns) | 15 | 1,<br>10 of top AF3,<br>10 of all AF3,<br>100 |
| 8OR8 **(2023)** | 1 (1 ns)<br>1 (10 ns)<br>2 (100ns) | 1 (1 ns)<br>1 (10 ns)<br>1 (100ns) | 18 | 1,<br>10 of top AF3,<br>10 of all AF3,<br>100 |

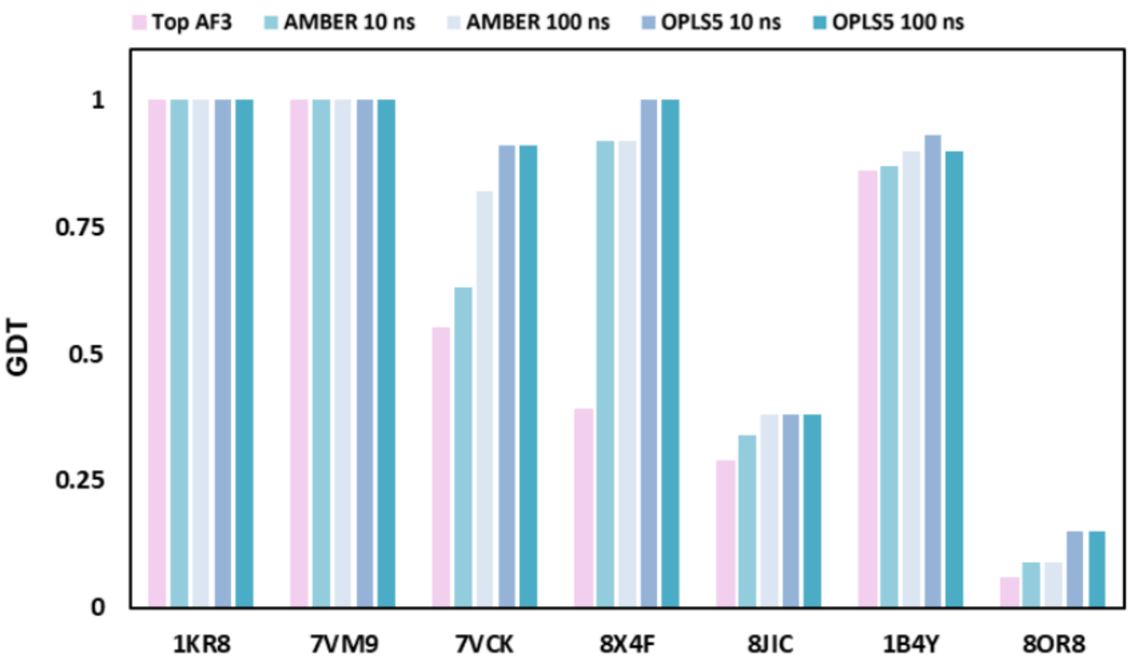


**Figure S1.** Best GDT score of the outputs from AF3 (pink), all results from AF3 each ran for 10 ns using ABMER99 (light teal) and OPLS5 (light lavender), and top results ran for 100 ns using ABMER99 (light navy) and OPLS5 (teal).

**Table S2.** Full list of the 36 cases studied, and their AF3 accuracy data before MD optimization. Errors in secondary structure predictions are listed. Bolded PDB IDs are cases reported in the manuscript. 28% (10 out of 36) of secondary structure was predicted correctly using SeqFold (MCC = 1), compared to 78% (28 out of 36) predicted correctly using AF3. 8% (3 out of 36) of the tertiary structure was predicted correctly using Fold/MMB (3Å cutoff GDT = 1.00), compared to 56% (20 out of 36) was predicted correctly using AF3.

| PBD (Year) | Len -gth | SeqFold MCC | Fold/MMB GDT | AF3 MCC | AF3 GDT | Structure | SeqFold Error | AF3 Error |
|---|---|---|---|---|---|---|---|---|
| **1KR8 (2002)** [23] | 7 | 0.47 | 1.00 | 1.00 | 1.00 | hairpin | 1 less pair | N/A |
| 1PQT (2003) [31] | 7 | 0.47 | 1.00 | 1.00 | 1.00 | hairpin | 1 less pair | N/A |
| 2K71 (2009) [32] | 8 | 0.58 | 1.00 | 1.00 | 0.86 | hairpin | 1 less pair | backbone error |

| | | | | | | | | |
|---|---|---|---|---|---|---|---|---|
| 5GWL (2016) [33] | 8 | 0 | 0.43 | 1.00 | 1.00 | 2 CCTG repeats | Predicted pair is wrong | N/A |
| 5GWQ (2016) [33] | 8 | 0.58 | 0.43 | 0.58 | 0.86 | 2 TTTA repeats | 1 pair predicted out of 2 | only predicted 1 out of 2 pairs |
| 6J37 (2019) [34] | 8 | 0.58 | 0.71 | 1.00 | 1.00 | minidumbbell structure of two CTTG repeats | Predicted pair is wrong | N/A |
| 6M0C (2020) [35] | 8 | -0.58 | 0 | 1.00 | 1.00 | minidumbbell formed by 5'-CTTG CATG-3' | Predicted pair is wrong | N/A |
| 2LO8 (2012) [36] | 10 | -0.17 | 0.33 | 1.00 | 1.00 | hairpin | Predicted pairs are wrong | N/A |
| 6IY5 (2020) [37] | 10 | 1 | 0 | 1.00 | 0.89 | minidumbbell formed by ATTCT repeats | N/A | backbone error |
| **7VM9 (2022)** [24] | 10 | 0.61 | 0 | 1.00 | 1.00 | minidumbbell formed with two regular CTTTG pentaloops | 1 out of 2 predicted | N/A |
| 7YF7 (2023) [38] | 10 | 1 | 0 | 1.00 | 0.89 | minidumbbell with 2 ATTTT repeat | N/A | backbone error |
| 1BJH (1997) [39] | 11 | 1 | 0.10 | 1.00 | 1.00 | hairpin | N/A | N/A |
| 2MKM (2014) | 11 | N/A | N/A | 1.00 | 1.00 | G triplex | No pairing predicted | N/A |

| | | | | | | | | |
|---|---|---|---|---|---|---|---|---|
| [40] | | | | | | | | |
| 2MKO (2014) [40] | 11 | N/A | N/A | 0.21 | 0.90 | G triplex | No pairing predicted | K+ was not in the AF3 input. there is a lot of ionic interactions |
| 5M1W (2017) [41] | 11 | N/A | N/A | 1.00 | 0.80 | G hairpin | No pairing predicted | backbone problem |
| 2LO5 (2012) [36] | 12 | 0.71 | 0.82 | 1.00 | 1.00 | hairpin | 3 out of 4 predicted | N/A |
| **7VCK (2022)** [29] | 12 | 1 | 0.45 | 1.00 | 0.55 | hairpin | N/A | Na+ not present. Backbone error |
| 1LA8 (2002) [42] | 13 | 1 | 0.17 | 1.00 | 1.00 | hairpin | N/A | N/A |
| 1P0U (2003) [43] | 13 | 1 | 0.08 | 1.00 | 1.00 | hairpin | N/A | N/A |
| 6R8E (2021) [44] | 14 | N/A | N/A | 0.19 | 0.62 | G hairpin | No pairing predicted | all Gs (G hairpin). but the pairs were shifted |
| **8X4F (2024)** [25] | 14 | 0.73 | 0.08 | 1.00 | 0.39 | hairpin | 1 missing pair | backbone problem |
| 1RDE (2003) [45] | 15 | N/A | N/A | 1.00 | 1.00 | G4 | No pairing predicted | N/A |

| | | | | | | | | |
|---|---|---|---|---|---|---|---|---|
| 2M8Y (2013) [46] | 15 | 1 | 0.07 | 1.00 | 1.00 | hairpin | N/A | N/A |
| 1D16 (1989) [30] | 16 | 1 | 0.13 | 1.00 | 1.00 | hairpin | N/A | N/A |
| 1AC7 (1997) [47] | 16 | 0.65 | 0.20 | 1.00 | 1.00 | hairpin | 1 missing pair | N/A |
| 7E5P (2021) [48] | 16 | -0.65 | 0 | 1.00 | 1.00 | G4 | No correct pairing | N/A |
| 1XUE (1997) [49] | 17 | 0.26 | 0 | 1.00 | 1.00 | hairpin | 2 pairs predicted correctly | N/A |
| 1ECU (2000) [50] | 19 | 1 | 0.44 | 1.00 | 1.00 | hairpin | N/A | N/A |
| **8JIC (2024)** [27] | 22 | -0.50 | 0 | 0.18 | 0.29 | G4 | No correct pairing | correct: anti//-chair, predicted: anti-// basket |
| 8RZX (2024) [51] | 24 | -0.35 | 0 | 1.00 | 0.17 | G4 | No correct pairing | unpaired region not predicted well |
| 1JVE (2002) [52] | 27 | 1 | 0.38 | 1.00 | 1.00 | hairpin | N/A | N/A |
| 1NGU (2003) [53] | 27 | 0.54 | 0.08 | 0.54 | 0.89 | hairpin | additional pairs predicted | additional pairs predicted |

| | | | | | | | | |
|---|---|---|---|---|---|---|---|---|
| | | | | | | | (identical to AF3 predictions) | |
| 7QB3 (2021) [54] | 28 | 0.56 | 0.04 | 0.19 | 0.07 | 2-way junction | Junction present but incorrect | junction not predicted correctly |
| **1B4Y (1999)** [26] | 30 | 0.62 | 0.07 | 1.00 | 0.86 | triple helix | Triplex missing only duplex | backbone problem |
| 8BM4 (2023) [55] | 32 | 0.29 | 0.03 | 0.09 | 0.23 | hairpin | Similar to AF3 predictions with few missing pairs | additional pairs and some backbone problems. both are hairpins though |
| **8OR8 (2023)** [28] | 35 | 0.70 | 0 | 0.25 | 0.06 | junction | Junction present but wrong | junction not predicted correctly |

**Table S3.** All 36 cases organized based on their motifs and their corresponding AF3-predicted tertiary structure accuracy.

| Name | 2nd structure | GDT |
|---|---|---|
| 1D16 | hairpin | 1 |
| 1XUE | hairpin | 1 |
| 1AC7 | hairpin | 1 |
| 1BJH | hairpin | 1 |
| 8BM4 | hairpin | 0.226 |
| 1ECU | hairpin | 1 |
| 1LA8 | hairpin | 1 |
| 1JVE | hairpin | 1 |
| 1KR8 | hairpin | 1 |

| | | |
|---|---|---|
| 1P0U | hairpin | 1 |
| 1NGU | hairpin | 0.885 |
| 1PQT | hairpin | 1 |
| 1RDE | hairpin | 1 |
| 2K71 | hairpin | 0.857 |
| 2LO8 | hairpin | 1 |
| 2LO5 | hairpin | 1 |
| 2M8Y | hairpin | 1 |
| 8X4F | hairpin | 0.385 |
| 6R8E | hairpin | 0.615 |
| 7VCK | hairpin | 0.545 |
| 5M1W | hairpin | 0.8 |
| 5GWL | repeat | 1 |
| 5GWQ | repeat | 0.857 |
| 6J37 | minidumbbell | 1 |
| 6IY5 | minidumbbell | 0.889 |
| 6M0C | minidumbbell | 1 |
| 7VM9 | minidumbbell | 1 |
| 7YF7 | minidumbbell | 0.889 |
| 1B4Y | triple helix | 0.862 |
| 2MKM | G triplex | 1 |
| 2MKO | G triplex | 0.9 |
| 7E5P | G4 | 1 |
| 8JIC | G4 | 0.286 |
| 8RZX | G4 | 0.174 |
| 7QB3 | junction | 0.57 |
| 8OR8 | junction | 0.51 |
